# Bacteriological and histopathological findings in cetaceans that stranded in the Philippines from 2017 to 2018

**DOI:** 10.1101/2020.11.30.403568

**Authors:** Marie Christine M. Obusan, Jamaica Ann A. Caras, Lara Sabrina L. Lumang, Erika Joyce S. Calderon, Ren Mark D. Villanueva, Cristina C. Salibay, Maria Auxilia T. Siringan, Windell L. Rivera, Joseph S. Masangkay, Lemnuel V. Aragones

**Affiliations:** Microbial Ecology of Terrestrial and Aquatic Systems, Institute of Biology, College of Science, University of the Philippines Diliman, Quezon City 1101, Philippines; Marine Mammal Research Stranding Laboratory, Institute of Environmental Science and Meteorology, College of Science, University of the Philippines Diliman, Quezon City 1101, Philippines; College of Science and Computer Studies, De La Salle University-Dasmariñas, City of Dasmariñas Cavite 4115, Philippines; Microbiological Research and Services Laboratory, Natural Sciences Research Institute, University of the Philippines Diliman, Quezon City 1101, Philippines; Pathogen-Host-Environment Interactions Research Laboratory, Institute of Biology, College of Science, University of the Philippines Diliman, Quezon City 1101, Philippines; College of Veterinary Medicine, University of the Philippines Los Baños, College, Batong Malake, Los Baños, Laguna 4031, Philippines

**Keywords:** cetaceans, histopathological assessment, lesions, antibiotic resistant bacteria, stranding events

## Abstract

The relatively high frequency of marine mammal stranding events in the Philippines provide many research opportunities. A select set of stranders (n=21) from 2017 to 2018 were sampled for bacteriology and histopathology. Pertinent tissues and bacteria were collected from eight species (i.e. *Feresa attenuata, Kogia breviceps, Globicephala macrorhynchus, Grampus griseus, Lagenodelphis hosei, Peponocephala electra, Stenella attenuata* and *Stenella longirostris)* and were subjected to histopathological examination and antibiotic resistance screening, respectively. Lesions that were observed in tissues of 19 cetaceans include congestion, hemorrhage, edema, hemosiderosis, glomerulopathy, Zenker’s necrosis, atrophy, atelectasis, and parasitic cysts. These lesions may be associated with factors possibly contributing to the death, debility, and stress of the animals during their strandings. On the other hand, the resistance profiles of 24 bacteria (belonging to genera *Escherichia, Enterobacter, Klebsiella, Proteus,* and *Shigella)* that were isolated from four cetaceans were determined using 18 antibiotics. All 24 isolates were resistant to at least one antibiotic class, and 79.17% were classified as multiple antibiotic resistant (MAR). The MAR index values of isolates ranged from 0.06 to 0.39 with all the isolates resistant to erythromycin (100%; n=24) and susceptible to imipenem, doripenem, ciprofloxacin, chloramphenicol, and gentamicin (100%; n=24). The resistance profiles of these bacteria can be used as basis for selecting antibiotics needed in the medical management of stranded cetaceans that need to be rehabilitated. Overall, the histopathological and bacteriological findings of the study demonstrate the challenges faced by cetacean species in the wild, such as but not limited to, biological pollution through land-sea movement of effluents, fisheries interactions, and anthropogenic activities.

## Introduction

The surveillance of wildlife health is part of an early warning system for detecting the emergence or resurgence of disease threats. In the case of cetacean populations in the Philippines, perhaps the most practical way of investigating their health is through their stranding events. A marine mammal is considered stranded when it runs aground, or in a helpless position such as when it is ill, weak, or simply lost [1]. While the event itself deserves attention, as it is not normal for any marine mammal to strand for no apparent reason, each stranded individual can give information on the abundance, distribution, health, and other ecological characteristics of its free-living counterparts [2], as well as threats faced by its population [3]. It is important that stranding events be responded as quickly as possible, since some stranded animals may quickly die depending on the size of the animal and extent of human intervention [4].

Biases exist in investigating the factors involved in cetacean strandings; easy-to-detect circumstances such as obvious injuries (especially those intentionally inflicted by humans) are likely to be more reported, whereas the role of diseases or parasites may be underestimated. The capacity to detect the presence of pathogens or parasites of stranded cetaceans depends on resources, such as the presence of a stranding network with the capability to respond to stranding events as well as availability of expertise for conducting necropsy and other protocols for case investigation. Nonetheless, whether or not a pathological condition is the underlying cause of a stranding, stranded animals are good representatives for monitoring wildlife health. Also, while live stranders provide good biological samples for laboratory analyses, a dead or decomposing carcass on the beach is just as useful in providing specimens and other information.

The available literature on bacteria that were isolated from marine mammals worldwide support the significance of investigating Gram-negative species and their antibiotic resistance or susceptibility. Antibiotic susceptibility patterns have been described for populations and individuals of Atlantic bottlenose dolphins, Pacific bottlenose dolphins, Risso’s dolphins, California sea lions, beluga whale, sea otters and pinnipeds [5–8]. Strains of zoonotic bacteria resistant to multiple antibiotics used for human and animal treatments were isolated from these animals, and some of those bacteria were recognized by the American Biological Safety Association (ABSA) as human pathogens. Associations between increased prevalence of antibiotic resistant bacteria in marine mammals and proximity to human activities were strongly suggested [5,7,9–11]. The antibiotic susceptibility profiles of bacteria isolated from cetaceans found in the Philippines where previously reported, wherein more than half of the bacteria (n=14) had single or multiple resistances to a selection of antibiotics [12].

On the other hand, histopathological assessments proved to be useful in determining probable causes of death or debility of stranded cetaceans worldwide [13–17]. Tissue lesions help confirm parasitic and bacterial infections, co-morbidities, physical injuries (e.g., brought about by fisheries or human interactions) and bioaccumulation of chemical compounds (e.g., persistent organic pollutants) in cetaceans [16, 18–22]. Histopathological assessment is a practical and informative tool that provides pathological evidence and reinforce the necropsy being done in dead cetaceans as part of the stranding response.

In this study, swab and tissue samples collected from cetaceans that stranded locally from February 2017-April 2018 were subjected to bacterial isolation (with subsequent antibiotic resistance screening) and histopathological assessment. Data on antibiotic resistant bacteria, parasites, and tissue lesions in cetaceans are valuable in evaluating the factors that may be associated with their local stranding events, observed to have increased in recent years [23–24]. Out of 29 confirmed species in the country, 28 species were reported to have stranded from 2005-2018 [24]. A yearly average of 105 cetaceans strandings occurred in the country from 2014 to 2018 [24]; 229 events were recorded by the Philippine Marine Mammal Stranding Network (PMMSN) in collaboration with the Bureau of Fisheries and Aquatic Resources (BFAR) from 2017 (n=121) to 2018 (n=108) involving 118 dead and 108 live (n=3 unknown) stranders.

## Materials and methods

All biological samples were collected in coordination with PMMSN and the Marine Mammal Research and Stranding Laboratory (MMRSL) of the Institute of Environmental Science and Meteorology (IESM), University of the Philippines, Diliman (UPD). The marine mammal stranding response and tissue collection is a nationwide effort which is part of the Memorandum of Agreement (MOA) between PMMSN and BFAR. Laboratory work was done at Microbial Ecology of Terrestrial and Aquatic Systems Laboratory (METAS) of the Institute of Biology, UPD.

### Sample collection

Cetaceans that stranded in the Philippines from February 2017 to April 2018 (Fig 1) were opportunistically sampled for tissues and swabs by veterinarians who were trained by PMMSN. Tissues were obtained during necropsy. Swabs were collected from routine (i.e., blowhole, anus, genital slit) and non-routine sites (e.g., organ sites and skin lesions suspected for infection) based on animal disposition and physical preservation [1]. Stranded cetaceans were characterized in terms of species, sex, age class, stranding type, stranding site, and stranding season (Table 1).

**Figure 1.**
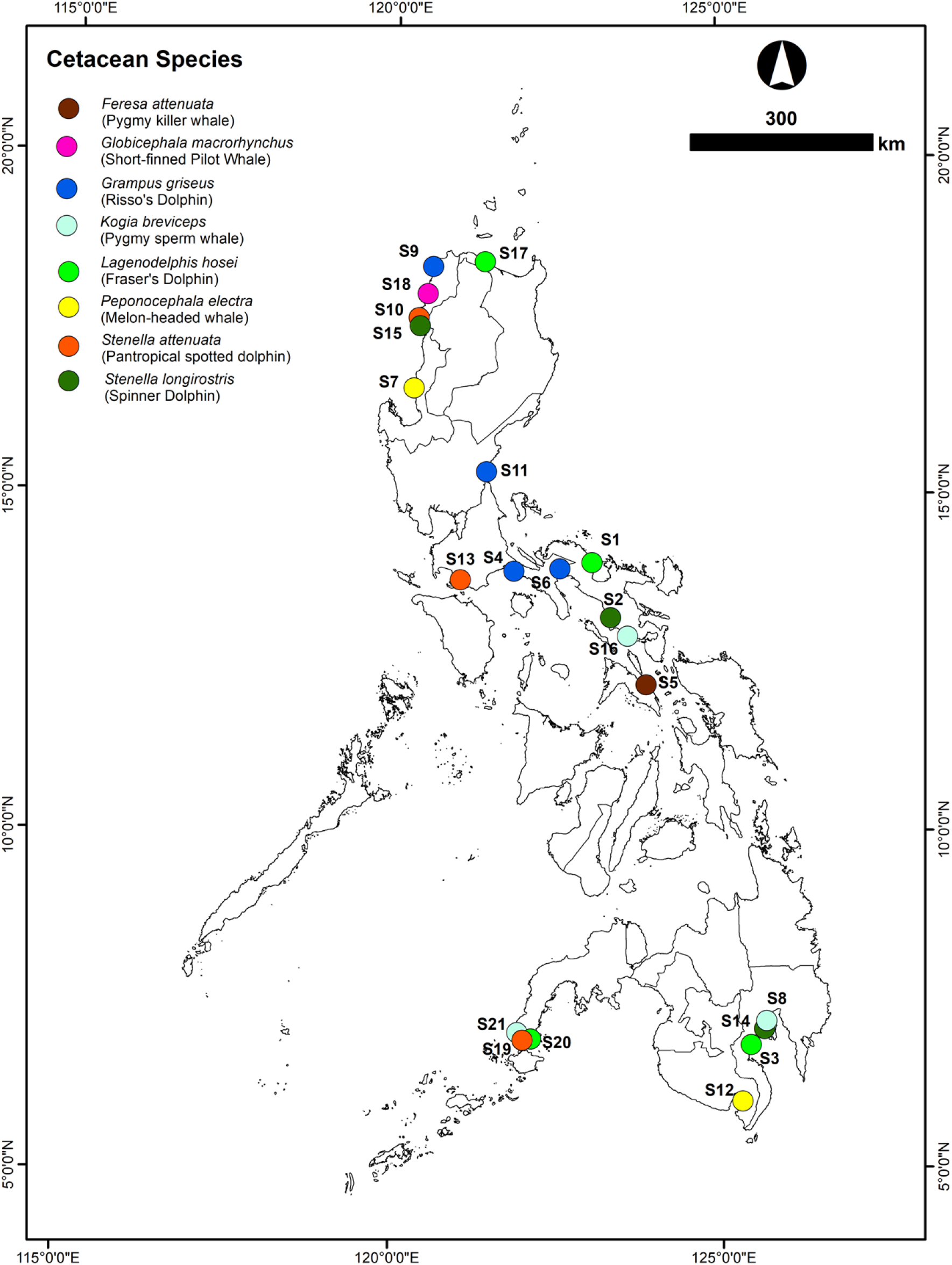
Sites of cetacean stranding events from February 2017-April 2018

**Table 1.** Stranded cetaceans responded for sampling in the Philippines during the year 2017 and 2018.

| Strander Code | PMMSN Code | Species | Region | Date | Stranding Type | Age Class | Condition | Sex |
| --- | --- | --- | --- | --- | --- | --- | --- | --- |
| S01 | Lh03R5270217 | Fraser's dolphin<br>*Lagenodelphis hosei* | V | 27-Feb-17 | Single | Adult | Alive | Male |
| S02 | Sl21R5040317 | Spinner dolphin<br>*Stenella longirostris* | V | 04-Mar-17 | Single | Adult | Alive | Female |
| S03 | Lh03R11010317 | Fraser's dolphin<br>*Lagenodelphis hosei* | XI | 09-Mar-17 | Single | Adult | Dead | Female |
| S04 | Gg04R4A290316 | Risso's dolphin<br>*Grampus griseus* | IV-A | 29-Mar-17 | Single | Subadult | Alive | Unknown |
| S05 | Fa02R5020517 | Pygmy killer whale<br>*Feresa attenuata* | V | 02-May-17 | Mass | Adult | Alive | Unknown |
| S06 | Gg15R5090517 | Risso's dolphin<br>*Grampus griseus* | V | 09-May-17 | Single | Adult | Alive | Unknown |
| S07 | Pe04R1300417 | melon-headed whale<br>*Peponocephala electra* | I | 30 April 2017 | Single | Unknown | Dead | Female |
| S08 | Kb07R11160517 | Pygmy sperm whale<br>*Kogia breviceps* | XI | 16-May-17 | Single | Adult | Alive | Male |
| S09 | Gg10R1150617 | Risso's dolphin<br>*Grampus griseus* | I | 15-Jun-17 | Single | Neonate | Alive | Female |
| S10 | Sa18R1210617 | pantropical spotted dolphin<br>*Stenella attenuata* | I | 21-Jun-17 | Single | Subadult | Dead | Female |
| S11 | Gg02R3230617 | Risso's dolphin<br>*Grampus griseus* | III | 23-Jun-17 | Single | Adult | Dead | Unknown |
| S12 | Pe06R12030717 | melon-headed whale<br>*Peponocephala electra* | XII | 03-Jul-17 | Single | Adult | Alive | Male |
| S13 | Sa03R4A280717 | pantropical spotted dolphin<br>*Stenella attenuata* | IV-A | 28-Jul-17 | Single | Subadult | Alive | Female |
| S14 | SI06R11310817 | spinner dolphin<br>*Stenella longirostris* | XI | 31-Aug-17 | Single | Subadult | Alive | Female |
| S15 | SI23R1300917 | spinner dolphin<br>*Stenella longirostris* | I | 30-Sep-17 | Single | Subadult | Dead | Female |
| S16 | Kb02R5091117 | pygmy sperm whale<br>*Kogia breviceps* | V | 09-Nov-17 | Single | Adult | Alive | Female |
| S17 | Lh04R2011217 | Fraser's dolphin<br>*Lagenodelphis hosei* | II | 01-Dec-17 | Single | Adult | Alive | Female |
| S18 | Gm11R151217 | short-finned pilot<br>whale<br>*Globicephala*<br>*macrorhynchus* | I | 05-Dec-17 | Single | Adult | Alive | Female |
| S19 | Sa03R9160118 | pantropical spotted<br>dolphin<br>*Stenella attenuata* | IX | 16-Jan-18 | Single | Adult | Alive | Female |
| S20 | Lh01R9170418 | Fraser's dolphin<br>*Lagenodelphis hosei* | IX | 17-Apr-18 | Single | Adult | Dead | Female |
| S21 | Kb01R9260418 | pygmy sperm whale<br>*Kogia breviceps* | IX | 27-Apr-18 | Single | Adult | Alive | Female |

### Histopathological assessment

Tissue samples (< 1 cm^3^ each) were preserved in 10% neutral buffered formalin, processed by paraffin-embedded technique, sectioned at 5 μm, and subjected to hematoxylin and eosin (H&E) staining. Tissue sectioning and H&E staining technique were performed at a private hospital Providence Hospital, Quezon City where tissue sections were stained with hematoxylin in water, dehydrated using a series of increasing concentrations of alcohol, and applied with eosin as a counterstain. Stained specimens were passed through xylol and toluol before mounting [25]. Using light microscopy technique, stained tissue samples were observed for the following pathologies: inflammation; fibrosis; granuloma lesions; edema; presence of parasite cysts (e.g., nematodes and protozoan parasites); endothelial damage (including endothelial deposits); presence of macrophages; granules; microthrombi formation; and hemorrhage.

### Bacterial isolation and antibiotic resistance screening

Swab samples in transport media (e.g., Amies) were stored at 4°C and were sent to the laboratory within 18-24 hours. Swabs were then enriched in Tryptic Soy Broth (TSB) for 18-24 hours at 37 °C. From the enriched media, inocula were streaked on MacConkey Agar (MCA). Morphologically distinct Gram-negative colonies were sub-cultured and purified.

Pure bacterial isolates were identified using 16S rRNA gene amplification. Bacterial DNA was extracted from the purified isolates using either the GF-1 Bacterial DNA Extraction Kit (Vivantis Technologies) following manufacturer’s instructions, or the Boil Lysis Method following Ahmed and Dablool (2017) [26]. The universal 16S rRNA bacterial gene was amplified from the DNA of isolates through polymerase chain reaction (PCR). The primers used for targeting the 16S rRNA gene were 27F (5’-AGAGTTTGATCCTGGCTCAG-3’) and 1541R (5’-AAGGAGGTGATCCANCCRCA-3’) [27–28]. The PCR reaction mix consisted of: dNTPs, MgCl2, *Taq* DNA polymerase, DNA template, forward and reverse primers, and nuclease-free water. The thermal cycler conditions were as follows: initial denaturation for 2 min at 95 °C, 30 cycles of denaturation for 30 s at 94 °C, annealing for 30 s at 55-60 °C, extension for 30 s at 72 °C, and final extension for 7 min at 72 °C. Positive controls (*E. coli* ATCC^®^ 25922) and blanks (DNA-free templates) were included. PCR products were subjected to agarose gel electrophoresis (AGE) to detect target DNA band. PCR products were then sent to a sequencing facility for DNA purification and sequencing. PreGap4 and Gap4 (Staden Package 2.0) were used to obtain the consensus sequences [29]. Sequence homologies were determined using NCBI BLASTn search and further analyses were done using BioEdit [30–31].

Kirby-Bauer Disk Diffusion Assay was performed to determine the sensitivity of the bacterial isolates to the following antibiotics: carbapenems (imipenem, meropenem, ertapenem, doripenem), penicillins (ampicillin), cephems (cephalothin, ceftriaxone, cefoxitin), fluoroquinolones (moxifloxacin, ciprofloxacin, ofloxacin), aminoglycosides (amikacin, gentamicin), tetracyclines (tetracycline, oxytetracycline), phenicols (chloramphenicol), folate pathway inhibitors (trimethoprim-sulfamethoxazole), and macrolides (erythromycin). These antibiotics were chosen based on (1) inclusion in the priority list of WHO for antibiotic resistance research; (2) known use in agriculture and aquaculture; (3) reported susceptibility profiles of bacteria isolated from marine animals worldwide; (4) use during rehabilitation of stranded marine mammals; and (5) known spectrum activity [5,32–35]. To ensure that only acquired resistances will be observed, antibiotics to which the bacterial isolates have intrinsic resistances were excluded in the assay. The reactions of the isolates to the antibiotics were described as Susceptible (S), Intermediate (I), or Resistant (R) based on Clinical and Laboratory Standards Institute (CLSI) M31-A2 (2002), M100-S24 (2014), and European Committee on Antimicrobial Susceptibility Testing (EUCAST) v 8.0 (2018). *E. coli* ATCC^®^ 25922 was used as the control [36–38]. Multiple Antibiotic Resistance (MAR) Index values were computed using the formula: (# of resistant antibiotics / total # of antibiotics tested) [39]. MAR indices greater than 0.2 were interpreted to come from sources where antibiotics are often used [32,39–40]. Also, MAR isolates were interpreted as those that are resistant to three or more antibiotic classes [41].

## Results

In this study, tissue samples and bacterial isolates were obtained from 21 stranded cetaceans representing eight species *(Feresa attenuata, Kogia breviceps, Globicephala macrorhynchus, Grampus griseus, Lagenodelphis hosei, Peponocephala electra, Stenella attenuata,* and *Stenella longirostris*). The stranding events of these cetaceans were responded in different regions of the Philippines, mostly in Region I, Region V, and Region IX, and the rest in Region II, Region III, Region IV-A, Region XI, and Region XII. Majority of the cetaceans (13) were female, while there are only three males. The sex of the remaining five cetaceans are unknown. For the age group, 15 were adults, five were sub-adults, and one was a neonate.

A total of 75 tissue samples from 19 out of 21 cetaceans were available for histopathological assessment (Table 2). The following lesions were observed in 44 tissue samples: congestion in 54.54% (n=24; brain, cardiac muscle, kidney, liver and lungs), hemorrhage in 13.63% (n=6; brain and kidney), edema in 13.63% (n=6; kidney, liver and lungs), hemosiderosis in 6.82% (n=3; brain and kidney), glomerulopathy in 20.45% (n=9; kidney), Zenker’s necrosis in 2.27% (n=1; skeletal muscle), atelectasis in 2.27% (n=1 lung), and parasitic cysts in 31.81%; *T. gondii* cysts in 11.36% (n=5; kidney and cardiac and skeletal muscles) *Sarcocystis* sp. cysts in 4.54% (n=2; skeletal muscle), and unidentified cysts in 15.91% (n=7; kidney, skeletal and cardiac muscles) (Fig 2).

**Table 2.** Histopathological findings in tissues of cetaceans that stranded in the Philippines from February to April 2018.

| Strander No. / Code | Species | Region | Date | Remarks in the Necropsy Report | Brain | Cardiac muscle | Kidney | Skeletal muscle | Liver | Lungs | Likely Cause of Stranding |
| --- | --- | --- | --- | --- | --- | --- | --- | --- | --- | --- | --- |
| S01 | *Lagenodelphis hosei* | V | 27-Feb-17 | unidentified parasites on eyes ; *P. delphini* cyst in the skeletal muscle | moderate congestion | NAL | NAL | unknown cyst | NTS | NTS |  |
| S02 | *Stenella longirostris* | V | 04-Mar-17 |  | moderate congestion | NAL | NAL | NAL | NTS | NTS |  |
| S03 | *Lagenodelphis hosei* | XI | 09-Mar-17 |  | moderate congestion | moderate congestion; *T. gondii* cysts | severe congestion; hemorrhage; edema | NAL | NTS | NTS | Shark attack |
| S04 | *Grampus griseus* | IV-A | 29-Mar-17 |  | NAL | severe congestion | severe congestion; | NAL | NTS | NTS |  |
| S05 | *Feresa attenuata* | V | 02-May-17 |  | moderate to severe congestion; hemorrhage; hemosiderosis | NTS | unidentified cysts; hemosiderosis | NAL | NTS | NTS |  |
| S06 | *Grampus griseus* | V | 09-May-17 |  | NAL | NAL | NTS | skeletal muscular atrophy; Zenker's necrosis; | NTS | NTS |  |
| S07 | *Peponocephala electra* | I | 30-April-17 |  | NTS | NAL | swollen glomerulus; hemosiderosis; | *T. gondii* cysts<br>nematode cyst | NTS | NTS |  |
| S08 | *Kogia breviceps* | XI | 16-May-17 | with encysted parasites in the muscle-blubber section, putative *P. delphini*; nematodes in the stomach | NAL | NAL | NTS | *Sarcocystis* cyst | NTS | NTS |  |
| S09 | *Grampus griseus* | I | 15-Jun-17 |  | NTS | severe congestion | NAL | nematode cyst | severe diffused hepatic sinusoidal congestion | severe congestion; focal pulmonary edema |  |
| S10 | *Stenella longirostris* | I | 21-Jun-17 |  | NAL | NAL | moderate congestion; hemorrhage; membranous glomerulopathy | *T. gondii* cysts | moderate to severe congestion | atelectasis | Acoustic trauma |
| S11 | *Grampus griseus* | III | 23-Jun-17 |  | NTS | NAL | NTS | NTS | NTS | NTS |  |
| S12 | *Peponocephala electra* | XII | 02-Jul-17 |  | NAL | NAL | membranous glomerulopathy | NAL | hepatic edema | pulmonary edema |  |
| S13 | *Stenella attenuata* | IV-A | 28-Jul-17 |  | NAL | NAL | glomerulopathy; edema | NTS | NTS | NTS |  |
| S14 | *Stenella attenuata* | XI | 31-Aug-17 | Abundant parasites in the stomach; external fresh wound due to cookie cutter shark bite | NTS | moderate congestion | glomerulopathy with lymphocytic aggregation; *T. gondii* cysts | *T. gondii* cyst<br>*Sarcocystis* cyst | NTS | NTS |  |
| S15 | *Stenella longirostris* | I | 30-Sep-17 |  | NAL | NAL | NTS | NAL | NAL | NAL | Acoustic trauma |
| S16 | *Kogia breviceps* | V | 09-Nov-17 | rope tied in peduncle area of fluke | severe congestion | unknown | hemorrhage; severe congestion | NTS | NTS | NTS |  |
| S17 | *Lagenodelphis hosei* | II | 01-Dec-17 |  | NTS | NTS | glomerulopathy; edema | unidentified cyst | NTS | NTS |  |
| S18 | *Globicephala macrorhynchus* | I | 05-Dec-17 |  | NAL | Moderate Congestion | hemorrhage; Glomerulopathy | unidentified cyst | NTS | NTS |  |
| S19 | *Stenella attenuata* | IX | 16-Jan-18 |  | NTS | severe congestion | hemorrhage; severe congestion; glomerulopathy | NTS | NTS | NTS |  |
| S20 | *Lagenodelphis hosei* | IX | 17-Apr-18 |  | severe congestion | severe congestion | severe congestion; glomerulopathy | NTS | NTS | NTS |  |
| S21 | *Kogia breviceps* | IX | 27-Apr-18 |  | NTS | moderate congestion | hemorrhage; glomerulopathy | NTS | NTS | NTS |  |

NAL, no apparent lesion; NTS, no tissue sample; Blank entries means no data

**Figure 2.**
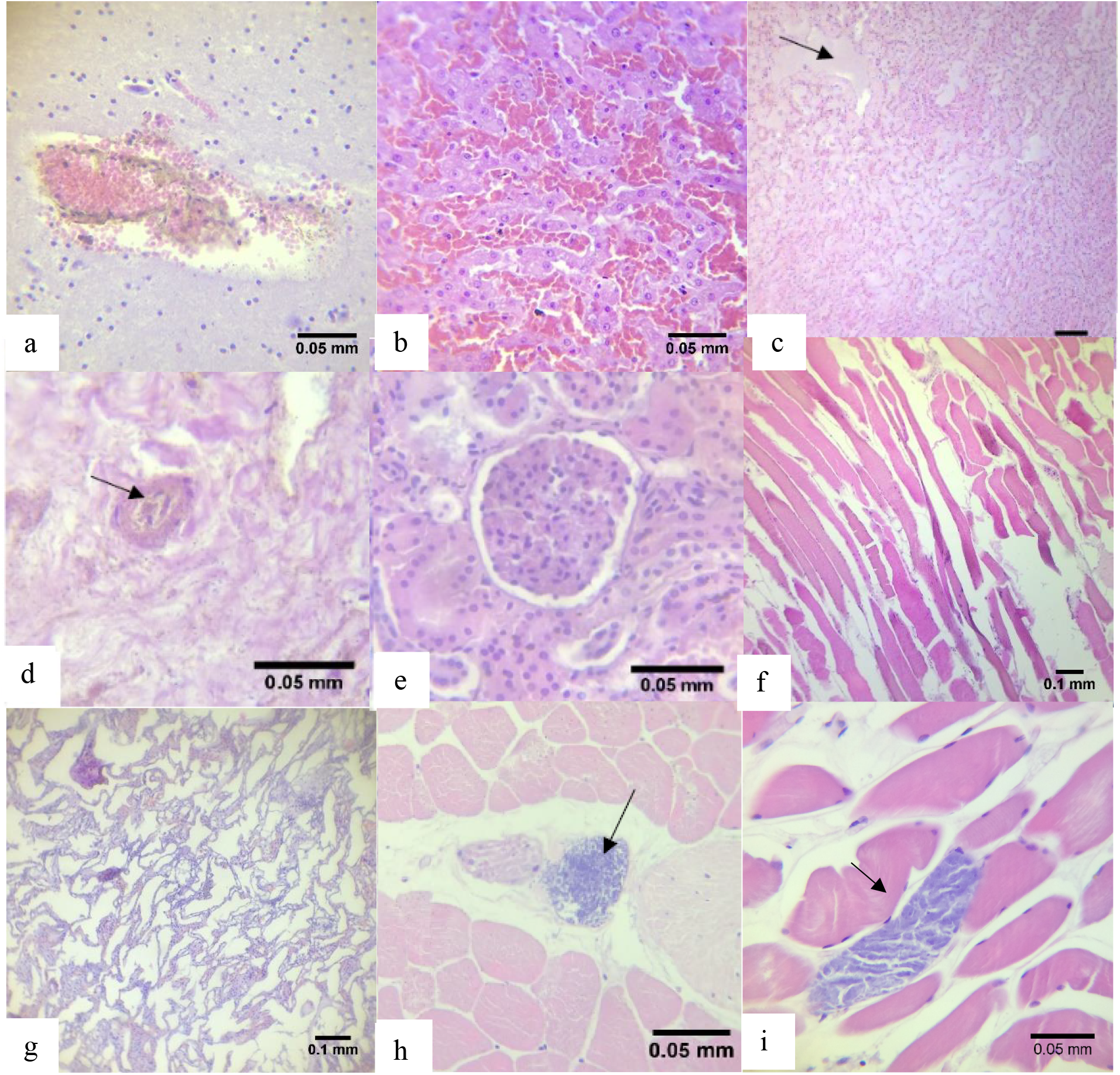
Some of the lesions observed in tissues. (a) hemorrhage in S03 kidney; (b) severe congestion in S11 liver; (c) edema in S12 liver; (d) Hemosiderosis characterized by the presence of brown granular pigments in S05 kidney; (e) glomerulopathy in S14 kidney; (f) Zenker’s necrosis characterized by hyaline degeneration, loss of striations, and muscle fiber waviness in S06 skeletal muscle; (g) atelectasis (collapsed alveoli) in S11 lungs; (h) *T. gondii* cyst in skeletal muscle of S14; and (i) *Sarcocystis* cyst in skeletal muscle of S08.

On the other hand, a total of 24 bacteria were isolated from four cetaceans (S12, S16, S17, S18). Based on 16S rRNA gene, these isolates were confirmed to have 98-100% sequence similarities to species belonging to the following genera: *Escherichia* (n=6), *Enterobacter* (n=8), *Klebsiella*(n=5), *Proteus* (n=4), and *Shigella* (n=1) (Table 3). These isolates were resistant to at least one antibiotic class tested, and 79.17% were classified as multiple antibiotic resistant (i.e., resistant to at least three antibiotic classes). The MAR index values of the isolates ranged from 0.06 to 0.39. (Fig 3).

**Table 3.**
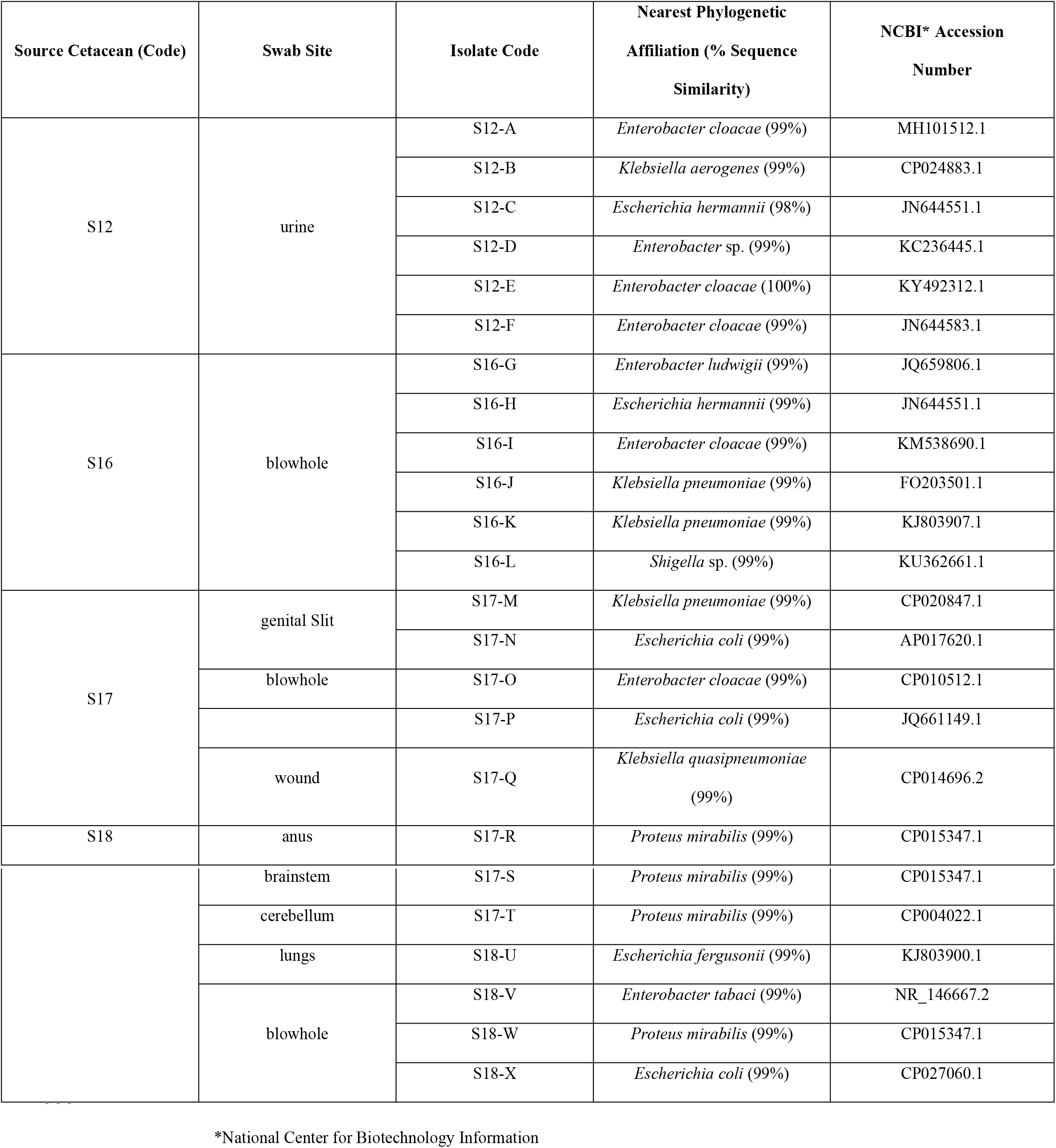
Genotypic identification of bacteria isolated from stranded cetaceans.

**Figure 3.**
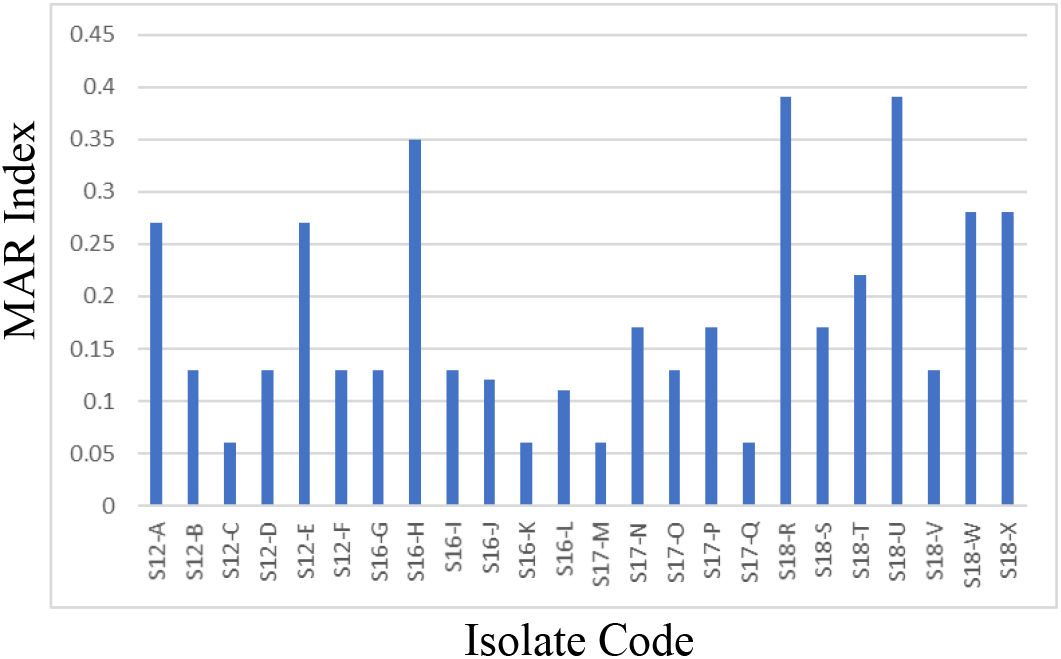
MAR (multiple antibiotic resistance) index values of Enterobacteriaceae isolated from cetaceans that straded in the Philippines from July to December 2017.

## Discussion

### Lesions in tissues of stranded cetaceans

Lesions in tissues of stranded cetaceans were associated with bycatch, trauma, parasitic and bacterial infections, and presence of persistent organic pollutants in stranded cetaceans [13–15,17,20,42–44]. A previous study in the Philippines involved the histopathological assessment of renal tissues which corroborated the results of molecular and culture methods for a suggested case of leptospirosis in a melon-headed whale *(Peponocephala electra)* [45]. However, the diagnosis of diseases through histopathological assessment of tissues of stranded marine mammals is often complex and difficult, involving many factors. Also, lesions may be caused by the stranding event itself and may mask pre-existing diseases.

When a cetacean strands, it is highly likely to have congestion in the liver and other organs due to the pressure from the weight of its body lying on the thorax as well as immotility preventing venous circulation [46]. The present study supports this, as congestion is the most observed type of lesion in hepatic and other organ tissues. Congestion in brain and kidneys of cetaceans has also been associated with acoustic trauma [47]. Acoustic trauma was suggested as the cause of some previously reported cetacean stranding events in the Philippines, possibly due to blast fishing activities near the stranding sites [23,48]. In addition, the stranding event itself can induce trauma and stress myopathy on the animal, causing congestion, hemorrhage, and skeletal and cardiac muscle degeneration such as in the case of Zenker’s necrosis [15,49–50]. These lesions were observed in 86% of tissues.

Glomerulopathy was observed in 20% of tissues with lesions. Comparably, membranous glomerulonephritis was a common finding among stranded cetaceans in Brazil [17]. This lesion was suggested to be associated with microbial infections or chronic exposure of cetaceans to metals such as cadmium, copper, and zinc, but this remains speculative due to the lack of toxicological analyses [44,51].

Parasites in cetaceans may predispose these animals to bacterial infections, cardiovascular complications, septicemia and other conditions, which are also frequently reported as probable causes of death during their stranding events [16,52–53]. Until this study, there was no histopathological evidence of coccidian cysts such as *T. gondii* and *Sarcocystis* sp. in muscles of locally-stranded cetaceans, with earlier reports on *T. gondii* detection using serological and molecular methods [45,54]. During the past two decades, coccidian infections have been detected in marine mammals that stranded along the coast of the northeastern Pacific Ocean [55–56]. These infections include encephalitis, myositis, hepatitis and myocarditis [57–58]. A better understanding of the biology, epidemiology, and pathogenesis of tissue-encysting coccidian organisms that parasitize marine mammals is needed to properly assess the risks and burden of protozoal disease in aquatic ecosystems [56–58]. In this study, cetaceans *Lagenodelphis hosei* (S03), *Peponocephala electra* (S07), *Stenella longirostris* (S10), and *Stenella attenuata* (S14) were found to have *T. gondii* cysts in their cardiac, kidney, and skeletal tissues while cetaceans *Kogia breviceps* (S08), and *Stenella attenuata* (S14) were found to have *Sarcocystis* spp. in their skeletal muscles. The transmission of these parasites is still poorly understood in marine mammals, although it is known that they are found in striated muscles of intermediate hosts [59–62]. The most likely modes of transmission of these parasites to aquatic animals are via ingestion of water-borne oocysts or sporocysts originating from sewage runoff or through infected prey [56-57,63-65].

In addition, the presence of *Phyllobothrium delphini* in the muscles and blubber of two sampled cetaceans was reported in the necropsy reports. This parasite has been documented in many cetacean species, commonly in the subcutaneous blubber with typical concentration in the perigenital region [66]. Siquier and Le Bas (2003) suggested that Fraser’s dolphins *(Lagenodelhis losei)* could act as intermediate or accidental hosts for *P. delphini*, and that definitive host infection could occur through predation. There is a need for more evidence to confirm the role of cetaceans in the life cycle of this parasite [67].

The consumption of muscles with these parasites is one of the major routes of transmission among hosts, including humans. This route of transmission is unlikely to involve cetaceans in the Philippines, as hunting and killing of marine mammals are prohibited under Section 4 of Republic Act 9147 (Wildlife Resources Conservation and Protection Act of the Philippines). Still, there were local reports of fishermen butchering cetaceans for food consumption (pers comm., BFAR Region V).

### Antibiotic resistant bacteria from stranded cetaceans

Overall, the bacterial isolates have resistances to carbapenems and third-generation cephalosporins. Enterobacteriaceae resistant to carbapenems and third-generation cephalosporins are considered a research priority for the discovery of new antibiotic agents [35]. As the “last line of defense” against multiple antibiotic resistant bacteria, the detection of carbapenem-resistant strains is a troubling point of concern as carbapenems are fourth-generation antibiotics recommended for critical Gram-negative infections [68]. To the best knowledge of the authors, only Greig et al. (2007) had so far used imipenem and meropenem for antibiotic susceptibility tests on bacteria isolated from cetaceans. Greig et al. reported imipenem-resistant *E. coli* in bottlenose dolphins, but all of their isolates were still susceptible to meropenem at the time [5].

All isolates were most resistant to erythromycin. The high frequency of resistance against this antibiotic is said to be due to acquired macrolide–lincosamide–streptogramin B (MLS) resistance genes, which is common among Enterobacteriaceae [69–70]. More than 50% of the isolates were also resistant to cephalothin, ampicillin, and moxifloxacin. It must be noted that the isolated *Klebsiella* spp., and *Escherichia* spp., bacterial species often reported as pathogenic to cetaceans, were resistant to erythromycin [71–72]. Similarly, high resistance to erythromycin, cephalothin, and ampicillin of *E. coli* isolated from bottlenose dolphins in Florida and South Carolina was reported [5–6]. Extra-intestinal pathogenic *E. coli* isolated from resident killer whales of San Juan Islands, Washington, were found to be resistant to aminoglycosides, sulfonamides, and tetracycline [73]. Resistances against cephalothin and ampicillin were also observed in bacteria isolated from dolphins, whales, and seals in the Northeastern United States Coast [32]. An overall high prevalence (88%) of resistance to at least one antibiotic was found among bacteria isolated from wild bottlenose dolphins in Florida, with highest resistances against erythromycin followed by ampicillin [6]. A previous study on antibiotic susceptibility patterns of bacteria isolated from stranded cetaceans in the Philippines reported the highest resistance (47%) to cefazolin [12].

The foregoing findings suggest that the mentioned antibiotics may not be good options for treating bacterial infections caused by Enterobacteriaceae in cetaceans. Based on the results of the study, the use of the following antibiotics must be considered in the medical management of stranded individuals (in order of decreasing potency): imipenem, doripenem, ciprofloxacin, chloramphenicol, gentamicin> meropenem, amikacin, ceftriaxone, trimethoprim-sulfamethoxazole> ertapenem> ofloxacin> tetracycline> cefoxitin> oxytetracycline> moxifloxacin> ampicillin, cephalothin> erythromycin. Susceptibilities to amikacin and gentamicin were also reported among bacteria isolated from marine mammals in Florida, South Carolina, and Northeastern US Coast [5,32].

The cetacean species sampled in this study generally inhabit deep waters, but their physiology entails a regular need to go up the surface to sequester oxygen from the air for breathing, thus exposing themselves to sewage outflows and other forms of pollution that eventually reach them from the nearby coast [74–76]. The presence of bacteria (and associated antibiotic resistances) in these cetaceans indicate biological pollution and presence of antibiotic resistance in their habitats [77–79]. In this study, 33.33% of the isolates from cetaceans had MAR indices greater than 0.2, suggesting that the isolates may have developed resistance from sources that the cetaceans were exposed to, such as bodies of water highly polluted with antibiotics, including domestic, industrial and hospital sewage outflows, water-treatment facilities, and the like [75–76]. As the use of antibiotics stems from anthropogenic activities, this implies the need to regulate and monitor the use and improper disposal of antibiotics to water bodies.

## Conclusion

Twenty-one cetaceans that stranded in different parts of the Philippines were sampled for histopathological assessment and bacterial isolation and antibiotic resistance screening. Histopathological findings showed congestion as the mostly observed lesion, followed by glomerulopathy, hemorrhage, and edema. In the absence of conclusive data on the specific causes of the mortality or morbidity of the cetaceans in relation to the stranding event, the observed lesions provide indications of possible involvement of acoustic trauma, stress, and chronic infections caused by microbial infections and exposure to chemical pollutants. On the other hand, bacteriological findings showed more than 50% of the isolated bacteria are multiple antibiotic resistant and that all of them are resistant to erythromycin and susceptible to imipenem, doripenem, ciprofloxacin, chloramphenicol, and gentamicin. While these information can serve as guide in the medical management of stranded cetaceans during rehabilitation, they also indicate the extent of antimicrobial resistance in the marine environment. As sentinels, cetaceans are demonstrating the threats faced by their populations in the wild, and monitoring their health through stranded representatives is a practical approach that can help improve conservation efforts. As local stranding network expands and veterinary and research expertise improve, more robust data from bacteriological and histopathological assessments of cetaceans are expected to be available in the coming years.

## Acknowledgments

We thank the Philippine Marine Mammal Stranding Network (PMMSN) and the Bureau of Fisheries and Aquatic Resources (BFAR) for the nationwide cetacean stranding response. Likewise, we thank Honey Leen M. Laggui for the preparation of Fig 1.

## Authors’ contributions

MCMO, JAAC, LSLL, EJSC, and RMDV performed the laboratory procedures. LVA led the cetacean stranding response with MCMO, JAAC, and RMDV. MCMO and LVA received funding for the study through project grants BIO-19-1-05 (UP-NSRI) and 171704SOS (UP-OVCRD) respectively. MCMO, LVA CCS, MATS, WLR, and JSM conceptualized the project, collated and analyzed all the data, and interpreted the results. All authors contributed to the writing and approved the final version of the manuscript.

